# Antibiotic Resistant *E. Coli* are Common in Waller Creek, an Urban Waterway Chronically Contaminated with Fecal Bacteria

**DOI:** 10.1101/2025.01.10.632463

**Authors:** Lena L. Olivera Perez, Stuart Reichler

## Abstract

The emergence of antibiotic-resistant bacteria has created a significant threat to global health. Some strains of *Escherichia coli* (*E. coli*), a common bacteria with both commensal and pathogenic strains, can be resistant to common antibiotics. This study investigates the prevalence of *E. coli* resistance to ampicillin, erythromycin, and ciprofloxacin in Waller Creek, an urban waterway in Austin, Texas with chronically high levels of fecal bacteria. Approximately 134 samples were collected from three different sites along Waller Creek and spiked with varying antibiotic concentrations. *E. coli* resistance to ampicillin and erythromycin was found to be up to 26.92% and 48.37% respectively, while resistance to ciprofloxacin was much lower ranging between 0%-2%. The highest levels of resistance were associated with dense urbanization, high levels of homelessness, and after heavy precipitation. These findings indicate the need for regular monitoring of antibiotic concentrations and antibiotic resistant fecal bacteria within urban waterways. This data is needed to identify potential human health risks and understand key factors that contribute to the quantity of antibiotic-resistant pathogens.

## Introduction

The creation of antibacterial therapy has become one of the most significant medical achievements of the twentieth century and is extensively utilized in modern medicine to save millions of lives. For example, prior to the antibiotic era, mortality rates for pneumonia caused by the bacterium *Streptococcus pneumoniae* was 40%, while the mortality rate for *Staphylococcus aureus* was as high as 80% (Friedman et al., 2016). But while antibiotics save lives, the extensive use of antibiotics in the healthcare and veterinary fields has facilitated the spread of antibiotic resistant bacteria, becoming a major health concern. Increases in antibiotic resistant bacteria have a number of clinical and economic drawbacks (Nij et al., 2021 and Oberle et al., 2012) and strains the healthcare system as a whole (Friedman et al., 2016 and Wei Bong et al., 2022).

*Escherichia coli* (*E. coli*) constitutes one of the most common commensal bacterial species present in the bowel flora (Erb et al., 2007). However, it is also a common pathogen causing urinary tract infections (UTI), pneumonia, bacteremia, peritonitis, diarrheal illnesses, among others (Mueller & Tainter, 2023). In 2020, 1 in 5 cases treated for a UTI caused by *E. coli* showed elevated resistance to standard antibiotics like ampicillin, co-trimoxazole, and fluoroquinolones (World Health Organization, 2023). Additionally, it is estimated that by 2050, the population who will die from antibiotic resistance will increase from 700,000 to 10 million per year globally. Each year in the United States, approximately 2.8 million individuals are infected with antibiotic resistant bacteria while 35,000 people die from these infections. In 2007 the European Centre for Disease Prevention and Control reported that 25,000 people died due to antibiotic resistant bacterial infections. The number of deaths increased to 33,000 in 2015 from 671,686 infected individuals. Death is not the only outcome of these infections. Many people suffer from lifelong disabilities increasing the amount of money that the government has to spend. For instance, more than 874,541 people received total disability-adjusted life years due to the impacts of antibiotic resistant infections in Europe (Nij et al., 2021).

The most common families of antibiotics are penicillins, tetracyclines, cephalosporins, fluoroquinolones, lincomycins, and macrolides (Anderson, 2023). For this project, the antibiotics studied were ampicillin, erythromycin, and ciprofloxacin, all from different families. Ampicillin is widely used for all types of simple infections while ciprofloxacin is only used for complicated infections and erythromycin can be used to treat susceptible infections (Anderson, 2023). The purpose of choosing these antibiotics was to determine the range of resistance from some of the most commonly used antibiotics.

A detectable percentage of antibiotics are excreted into aquatic environments through urine and feces by coupling with polar molecules or non-metabolized forms. Antibiotic residues in wastewater constitute emergent contaminants of water systems in urban environments (Oberle et al., 2012). Ampicillin resistance in *E. coli* has become widespread in humans and the environment. A 2003 study of wastewater in Austria found 18% of *E. coli* resistant to ampicillin (Reinthaler et al., 2003). Another study performed around a wastewater treatment plant near a medical center showed that 38% of the total samples were resistant to ampicillin (Oberle et al., 2012). Additionally, a multi-sectional study published in 2021 synthesized studies conducted from 1989 to 2019 in urban areas and 17% of 10,618 water samples were resistant to ciprofloxacin (Nij et al., 2021). A report from 2023 showed that *E. coli* resistance to ciprofloxacin increased steadily from 14.2% to 19.8% from the years 2015 to 2021, even as prescriptions for ciprofloxacin decreased three-fold during the same period (Tchesnokova et al., 2023). Moreover, a longitudinal comparison study of antibiotic resistance in 2016 showed that *E. coli* resistance to erythromycin ranged from 27.2% to 45.3% (Seidman et al., 2016).

This is a preliminary study with the goal of determining the levels of antibiotic resistance in *E. coli* to different antibiotics and in different conditions in an urbanized creek with chronically high fecal bacteria levels. While finding antibiotic resistant *E. coli* is not necessarily cause for alarm, it does indicate that further monitoring would be prudent.

## Materials and Methods

### Sample Collection

Sample collection took place at three sites along Waller Creek in Austin, TX: Creekside (CS), 43^rd^ street (FT), and 7th Street (SS). These locations were selected because previous monitoring showed them to have high levels of *E. coli* (data not shown).

Water was collected as per Riedel et al. (2015) in 500 mL amber bottles (previously rinsed with 10% HCl) between 8:00 AM and 11:00 AM in areas of flowing water. Each amber bottle was rinsed three times downstream before they were fully submerged upstream to collect the sample. The bottles were placed at 4°C until sample preparation started which was no more than 4 hours after collection.

### Impervious Cover calculations

Impervious cover calculations were estimated using iTree Canopy (https://canopy.itreetools.org/). A 1 square kilometer area upstream of the sample site was used to calculate impervious cover using 100 points for each location. The iTree Canopy categories impervious roads, impervious buildings, and impervious others were added together to determine the estimated impervious cover for each site.

### Antibiotic Stock Solutions

Stock solutions were created for three antibiotics: ampicillin (100 mg/mL), erythromycin (50 mg/mL resuspended in DMSO), and ciprofloxacin (10 mg/mL). Each stock solution was filter sterilized (0.22 μm) and stored at -20°C.

### Sample Preparation and E. coli Quantification

For each sample 50 mL of creek water, 50 mL of DI water, the appropriate amount of antibiotic stock solution, and the Colilert reagent were added to a 100 mL graduated cylinder. The reagent utilizes proprietary Defined Substrate Technology in order to detect coliforms and *E. coli* (IDEXX, 2024). Control samples had no antibiotic stock solution added to them. Since erythromycin has low solubility in water, DMSO was used for the stock solution, and samples with erythromycin had two control groups in order to see if there was any difference when DMSO was used as solvent compared to water. The sample was poured into the Quanti-Tray/2000 and sealed by the Quanti-Tray Sealer. Each concentration had three replicates and the procedure was repeated for each of them. After all the trays were sealed, they were placed in an incubator at 35°C for 24 ±2 hours.

After incubation the trays were removed and the Most Probable Number (MPN) of *E. coli* was calculated following the procedure from IDEXX, 2024. The MPN of *E. coli* was then multiplied by two to account for the dilution factor.

## Results

Samples were collected on 7 different days and 3 sample locations for a total of 134 water samples with 57 samples spiked with ampicillin, 26 samples with erythromycin, and 51 samples with ciprofloxacin (Table 1).

**Table 1.**
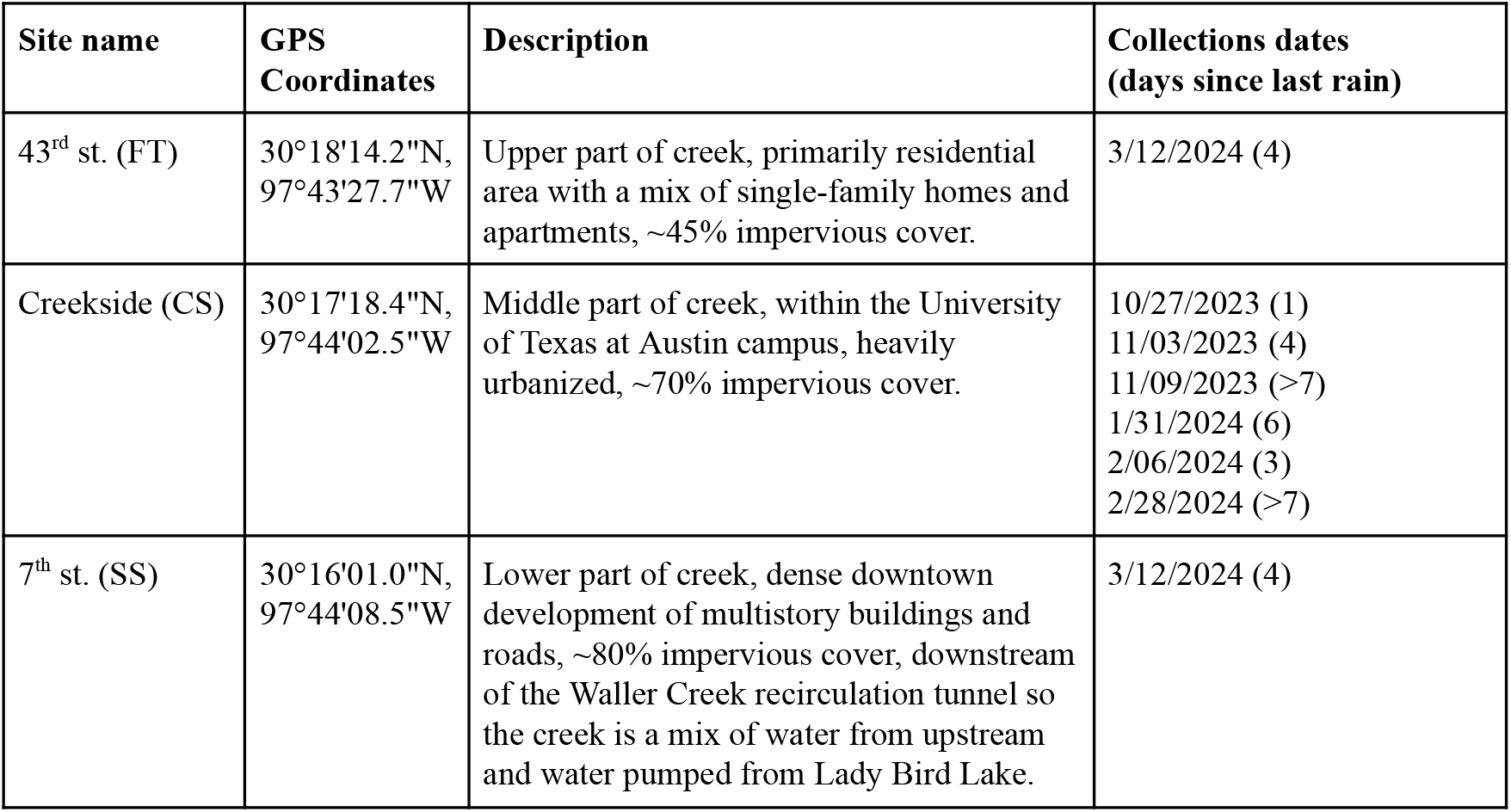
Waller Creek Sample site descriptions including location, impervious cover, and days since last rain.

| Site name | GPS Coordinates | Description | Collections dates (days since last rain) |
| --- | --- | --- | --- |
| 43<sup>rd</sup> st. (FT) | 30°18'14.2"N,<br>97°43'27.7"W | Upper part of creek, primarily residential area with a mix of single-family homes and apartments, ~45% impervious cover. | 3/12/2024 (4) |
| Creekside (CS) | 30°17'18.4"N,<br>97°44'02.5"W | Middle part of creek, within the University of Texas at Austin campus, heavily urbanized, ~70% impervious cover. | 10/27/2023 (1)<br>11/03/2023 (4)<br>11/09/2023 (>7)<br>1/31/2024 (6)<br>2/06/2024 (3)<br>2/28/2024 (>7) |
| 7<sup>th</sup> st. (SS) | 30°16'01.0"N,<br>97°44'08.5"W | Lower part of creek, dense downtown development of multistory buildings and roads, ~80% impervious cover, downstream of the Waller Creek recirculation tunnel so the creek is a mix of water from upstream and water pumped from Lady Bird Lake. | 3/12/2024 (4) |

### Ampicillin Resistance

Figure 1a shows the average MPN of *E. coli* present in each sample after incubation by water sample collection date and location for ampicillin. On October 27, 2023 (light green), samples were collected at the Creekside site and, as can be seen, the average MPN of *E. coli* measured was 1,668 MPN/100 mL for the control group (n = 2). On the same day, samples with a concentration of 20 μg/mL (n = 2), 50 μg/mL (n = 2), and 200 μg/mL (n = 2) of ampicillin averaged 449 MPN/100 mL, 246 MPN/100 mL, and 62 MPN/100 mL respectively. Similarly, on November 3, 2023 (light orange), Creekside was visited again and the average MPN of *E. coli* present in the control group (n = 3) was lower at 819 MPN/100 mL while the MPN measured in samples with a concentration of 20 μg/mL (n = 3), 50 μg/mL (n = 3), and 200 μg/mL (n = 3) of ampicillin averaged 118 MPN/100 mL, 10 MPN/100 mL, 6 MPN/100 mL respectively. Six days later, on November 9, 2023 (light blue), at the same sample site, the average *E. coli* present in the control sample (n = 3) was 653 MPN/100 mL and for samples with a concentration of 20 μg/mL (n = 3), 50 μg/mL (n = 3), and 200 μg/mL (n = 3) of ampicillin, the average MPN of *E. coli* was 13.3 MPN/100 mL, 10 MPN/100 mL, and 0 MPN/100 mL respectively. The collection site at 43^rd^ street was visited March 12, 2024 (dark blue), and it showed an *E. coli* count for the control group (n = 3) of 308 MPN/100 mL while concentrations of 20 μg/mL (n = 3), 50 μg/mL (n = 3), and 200 μg/mL (n = 3) of ampicillin averaged 20 MPN/100 mL, 9 MPN/100 mL and 2 MPN/100 mL respectively. On the same day, the collection site at 7^th^ street (purple), showed an average *E. coli* MPN of 1,028 MPN/100 mL for the control group (n = 3). For concentrations of 20 μg/mL (n = 3), 50 μg/mL (n = 3), and 200 μg/mL (n = 3) of ampicillin, the average count was 71 MPN/100 mL, 42 MPN/100 mL, and 4 MPN/100 mL respectively.

**Figure 1a.**
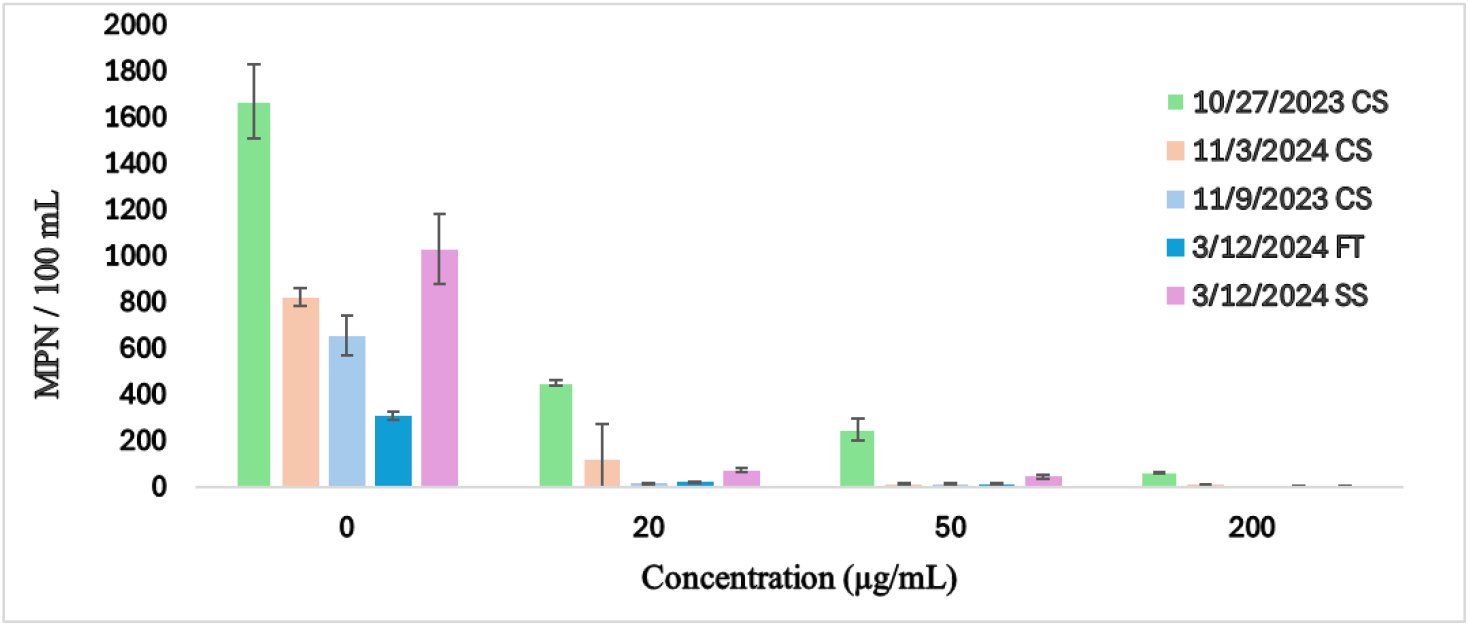
Most Probable Number of *E. coli*/100mL present after addition of ampicillin.

Since the number of *E. coli* for the no antibiotic control changed by collection date and location, figure 1b shows the percent of *E. coli* remaining as the concentration of ampicillin increases. Each control group is represented as 100% of *E. coli* measured for that particular day. On October 27, 2023 (light green), samples with a concentration of 20 μg/mL, 50 μg/mL, 100 μg/mL and 200 μg/mL of ampicillin represented 27%, 15%, 13%, and 4%, respectively, of the MPN of *E. coli* present in the control group. On November 3, 2023 (light orange), samples with a concentration of 20 μg/mL of ampicillin showed 14% of the *E. coli* MPN compared to what was measured in the control group. Furthermore, samples with a concentration of 50 μg/mL and 200 μg/mL of ampicillin were 1% and 1% of the total count. On November 9, 2023 (light blue), samples with a concentration of 20 μg/mL and 50 μg/mL of ampicillin represented 2% and 2% respectively of the average MPN in the control group. Consequently, on March 12, 2024 at 43rd Street (dark blue), samples with a concentration of 20 μg/mL, 50 μg/mL, and 200 μg/mL of ampicillin represented 6%, 3% and 1% (respectively) of the total MPN of *E. coli* found at the site. On the same day, but at 7^th^ street, samples with a concentration of 20 μg/mL, 50 μg/mL, and 200 μg/mL of ampicillin were 7%, 4% and 0.4%, respectively, of the MPN measured in the control group.

**Figure 1b.**
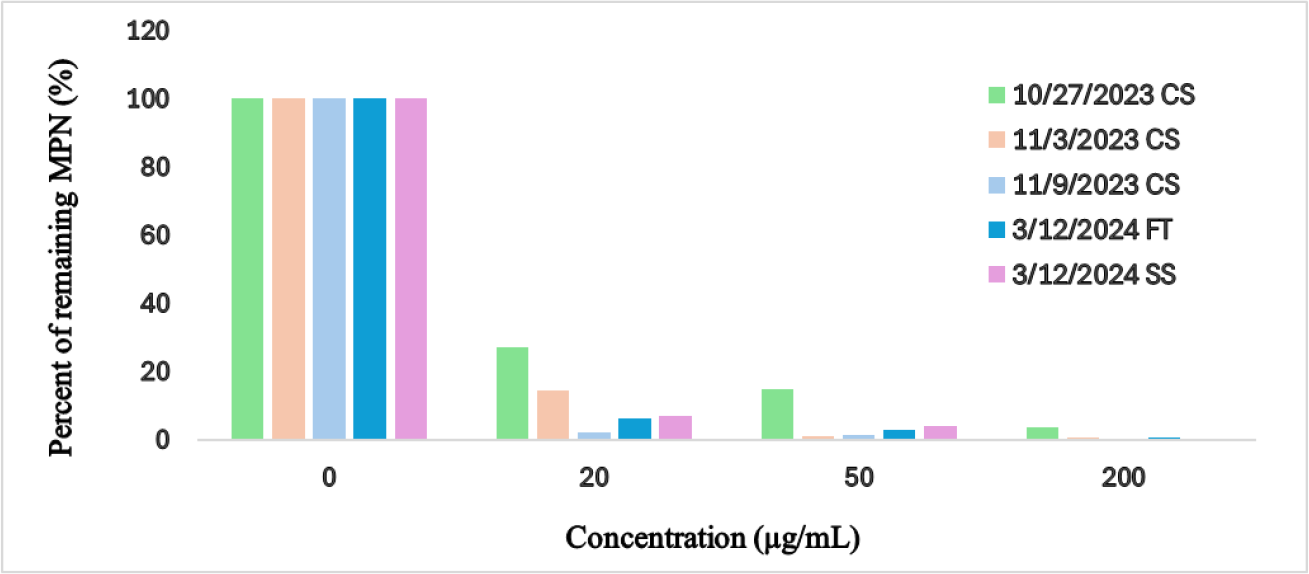
Proportion of remaining *E. coli* present after addition of ampicillin.

A paired t-test was performed to compare the average *E. coli* count of the control group to the average *E. coli* count of each concentration for a particular day in order to determine if the decrease of *E. coli* was statistically significant. As noted in Table 2, all P-values are less than 0.05 which indicates that the differences were statistically significant.

**Table 2.** Paired t-test results for ampicillin resistance.

| Concentration<br>($\mu\text{g/mL}$) | 10/27/2023<br>CS | 11/03/2023<br>CS | 11/09/2023<br>CS | 03/12/2024<br>FT | 03/12/2024<br>SS |
| --- | --- | --- | --- | --- | --- |
| 0 vs. 20 | 0.02 | 0.003 | 0.0005 | 0.0003 | 0.0009 |
| 0 vs. 50 | 0.01 | 0.0007 | 0.0005 | 0.0002 | 0.0008 |
| 0 vs. 200 | 0.01 | 0.0007 | 0.0007 | 0.0002 | 0.0007 |

### Erythromycin Resistance

Figure 2a represents the average *E. coli* count present in each erythromycin sample after incubation by collection day and location. On February 28, 2024 (dark orange), the average MPN of *E. coli* present in the control sample (n = 3) and the DMSO control sample was 326 MPN/100 mL. Consequently, the average MPN of *E. coli* for samples with a concentration of 10 μg/mL (n = 3) and 20 μg/mL (n = 3) of erythromycin was 158 MPN/100 mL and 12 MPN/100 mL respectively. On March 12, 2024 (dark blue), at the 43^rd^ street collection site, the *E. coli* count for the control group (n = 3) and the control DMSO samples averaged 321 MPN/100 mL and 311 MPN/100 mL respectively while the average MPN of *E. coli* seen in samples with a concentration of 10 μg/mL (n = 3) and 20 μg/mL (n = 3) of erythromycin was 62 MPN/100 mL and 5 MPN/100 mL respectively. On the same day but at 7^th^ street, the average *E. coli* count for the control group and the DMSO control group (n = 3) was 1,048 MPN/100 mL and 1,039 MPN/100 mL, respectively. Finally, the average MPN of *E. coli* present in samples with a concentration of 10 μg/mL (n = 3) and 20 μg/mL (n = 3) of erythromycin was 880 MPN/100 mL and 53 MPN/100 mL, respectively.

**Figure 2a.**
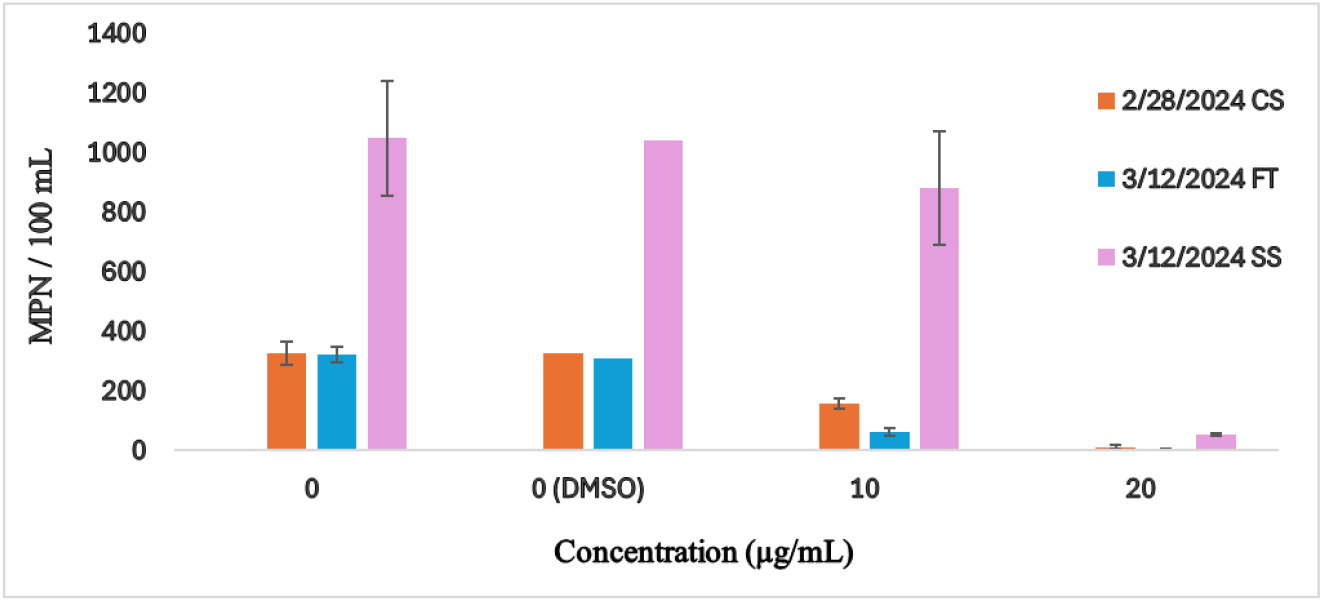
Most Probable Number of *E. coli*/100mL present after addition of erythromycin.

Since the number of *E. coli* for the no antibiotic control changed by collection date and location, figure 2b shows the percentage of remaining *E. coli* present in the samples as the concentration of erythromycin increased. The control groups of each day represent 100% of the *E. coli* present in each sample. The DMSO control samples showed no statistical difference compared to the water only control. On February 28, 2024 (dark orange), the samples with concentrations of 10 μg/mL and 20 μg/mL of erythromycin represented 48% and 4% respectively of the total *E. coli* found in the control. Consequently, on March 12, 2024, on 43^rd^ Street (dark blue), the samples with concentrations of 10 μg/mL and 20 μg/mL of erythromycin were 19% and 2% of the total *E. coli* count. On the same day but on 7^th^ street (purple), samples with 10 μg/mL and 20 μg/mL of erythromycin represented 84% and 5% of the total count of *E. coli*.

**Figure 2b.**
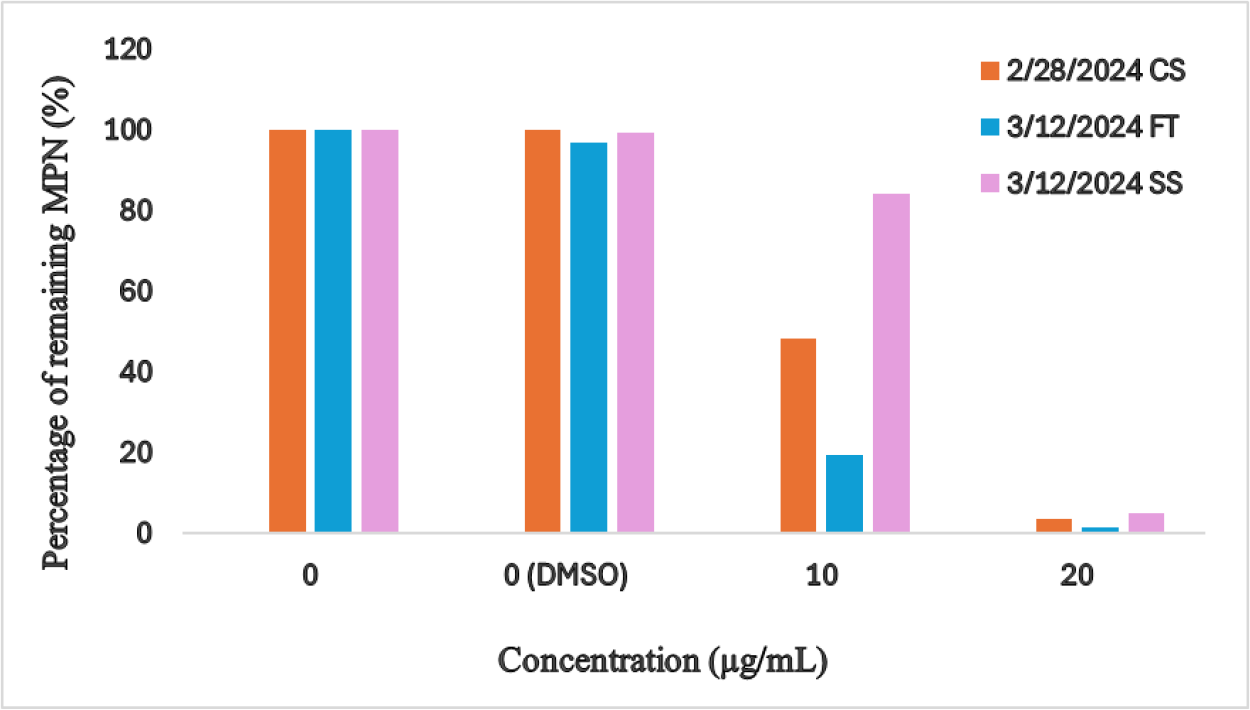
Proportion of *E. coli* present after the addition of erythromycin.

The control group and the DMSO control group were the same, and the control without DMSO was used as the control for the statistical analysis.. In order to identify if the difference between control and spiked samples were statistically significant, a paired t-test was performed. As can be seen in Table 3, the differences between the control group and the samples with concentration 20 μg/mL of erythromycin were statistically significant for all days. The differences between the control group and samples with 10 μg/mL of erythromycin were statistically significant for most days except March 12, 2024 at the collection site located at 7^th^ street.

**Table 3.** Paired t-test results for erythromycin resistance.

| Concentration<br>(µg/mL) | 02/08/2024<br>CS | 02/28/2024<br>CS | 03/12/2024<br>FT | 03/12/2024<br>SS |
| --- | --- | --- | --- | --- |
| 0 vs. 10 | 0.006 | 0.006 | 0.0002 | 0.4 |
| 0 vs. 20 | 0.0004 | 0.0004 | 0.00006 | 0.01 |

### Ciprofloxacin Resistance

Figure 3a shows the average *E. coli* present in the samples with ciprofloxacin by collection dates and location. On January 31, 2024 (light blue), at the Creekside location, the average *E. coli* count was 185 MPN/100 mL for the control groups (n = 3) while the count for samples with concentrations of 1μg/mL (n = 3) and 5 μg/mL (n = 3) of ciprofloxacin was 3 MPN/100 mL and 3 MPN/100 mL. On the same site but on February 6, 2024 (green), the *E. coli* count averaged 546 MPN/100 mL for the control samples (n = 3) and 2 MPN/100 mL (n = 3) for the samples with a concentration of 0.1 μg/mL (n = 3) of ciprofloxacin. Samples with a concentration of 1μg/mL (n = 3) and 5 μg/mL (n = 3) of ciprofloxacin had no *E. coli*. On February 28, 2024 (orange), at the same collection site, the average number of *E. coli* was 360 MPN/100 mL while the average count for samples with concentrations of 0.1μg/mL (n = 3) and 1 μg/mL (n = 3) of ciprofloxacin was 2 MPN/100 mL and 0.7 MPN/100 mL, respectively. Similarly, samples with a concentration of 5 μg/mL (n = 9) of ciprofloxacin did not have any surviving *E. coli*. At 43^rd^ Street (dark blue), the average count for *E. coli* was 331 MPN/100 mL for the control group (n = 3) while samples with concentrations of 0.1 μg/mL (n = 3) and 1 μg/mL (n = 3) of ciprofloxacin had 3 MPN/100 mL and 3 MPN/100 mL respectively. At 7^th^ Street (purple), the control sample (n = 3) had an average *E. coli* count of 957 MPN/100 mL while samples with concentrations of 0.1 μg/mL (n = 3) and 1 μg/mL (n = 3) had an average count of 8 MPN/100 mL and 8 MPN/100 mL respectively.

**Figure 3a.**
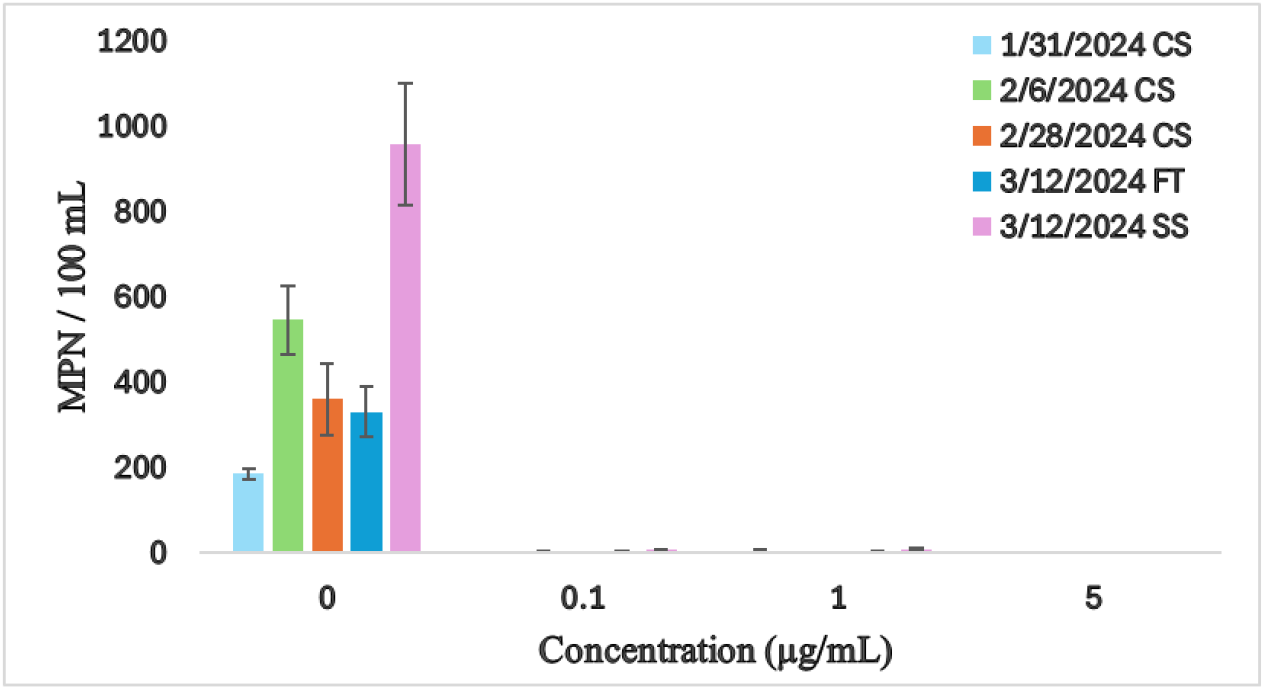
Most Probable Number of *E. coli*/100mL present after addition of ciprofloxacin.

Since the number of *E. coli* for the no antibiotic control changed by collection date and location, figure 3b represents the percentage of *E. coli* that were still present in the samples after the concentration of ciprofloxacin was increased. The control is represented as 100% of the total number of *E. coli* present. On January 31, 2024 (light blue), samples with a concentration of 1 μg/mL and 5 μg/mL of ciprofloxacin represented 2% and 1% of the total *E. coli* count while, on February 6, 2024 (green), samples with a concentration of 0.1 μg/mL of ciprofloxacin were 0.4% of the total number of *E. coli* that day. On February 28, 2024 (orange), the samples at Creekside with a concentration of 0.1 μg/mL and 1 μg/mL of ciprofloxacin represented 0.6% and 0.2% of the control samples respectively. At the 43^rd^ Street site (dark blue), samples with a concentration of 0.1 μg/mL and 1 μg/mL of ciprofloxacin represented 1% and 1% of the control group respectively while the same concentrations represented 1% and 1% respectively at 7^th^ Street (purple).

**Figure 3b.**
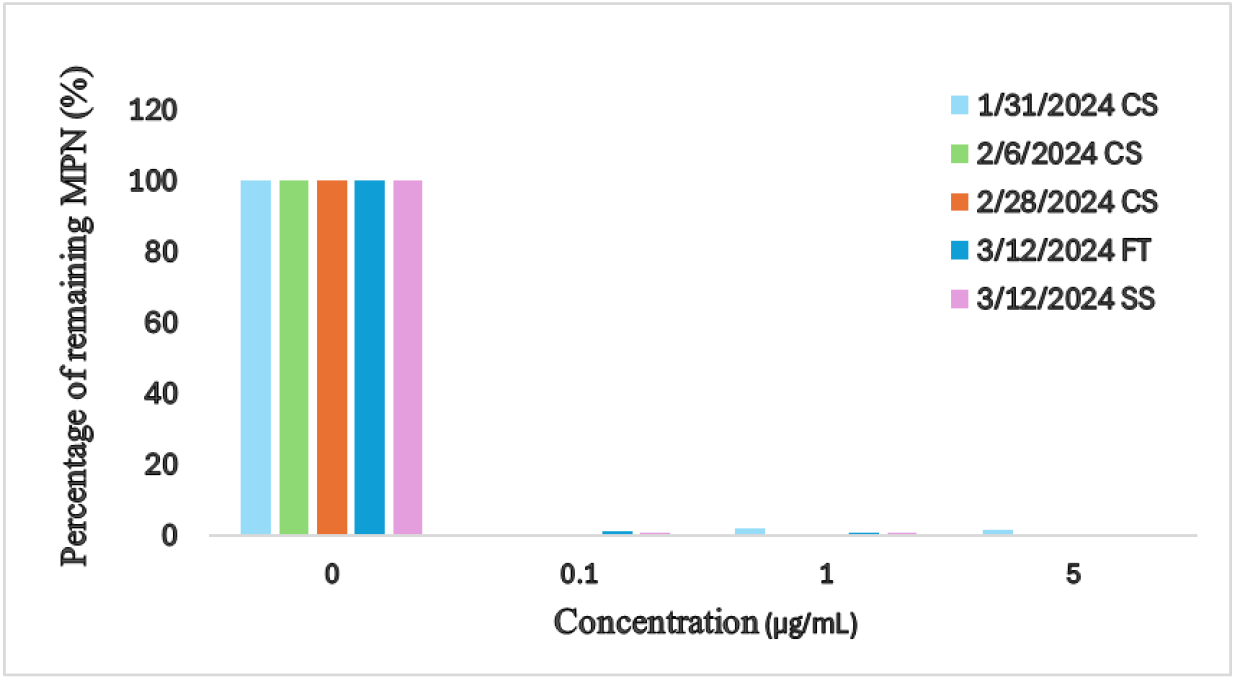
Proportion of remaining *E. coli* present after addition of ciprofloxacin.

A paired t-test was performed to determine if the difference between the samples with ciprofloxacin and the control were statistically significant. As can be noted in Table 4, all of the calculated P-values are less than 0.05 thus, the difference between the control and spiked samples was statistically significant.

**Table 4.** Paired t-test results for ciprofloxacin resistance.

| Concentrations<br>(µg/mL) | 01/31/2024<br>CS | 02/06/2024<br>CS | 02/28/2024<br>CS | 03/12/2024<br>FT | 03/12/2024<br>SS |
| --- | --- | --- | --- | --- | --- |
| 0 vs. 0.1 | - | 0.0007 | 0.004 | 0.001 | 0.0007 |
| 0 vs. 1 | 0.0003 | 0.0007 | 0.004 | 0.001 | 0.0007 |
| 0 vs. 5 | 0.0004 | 0.0007 | 0.004 | - | - |

## Discussion

This project’s goal was to determine if antibiotic resistant bacteria are present in an urban creek with a known history of high fecal bacteria levels, and if so, the level of *E. coli* resistance to common antibiotics focusing on ampicillin, erythromycin, and ciprofloxacin. The results indicate that *E. coli* in Waller Creek have a moderate resistance to ampicillin and erythromycin while very little resistance was observed for ciprofloxacin.

The level of ampicillin resistance found in Waller Creek (up to 27%) is in a similar range found by Reinthaler et al. (2003) and Oberle et al. (2012), 18% and 38% respectively. The percent of resistance is also comparable to a study conducted in the Mur River and Drava River in Austria, which showed moderate resistance of *E. coli* to ampicillin ranging from 14% to 16% (Skof et al., 2024). Similarly, another study conducted in the Qiantang River and the Dongdiao Stream in Hangzhou City, China found 29% *E. coli* resistance to ampicillin (Chen et al., 2017). One of the differences between these four studies was the type of water being tested: the studies of Reinthaler et al. (2003) and Oberle et al. (2012) were conducted in sludge and sewage while the research of Chen et al. (2017) and Skof et al.(2024) worked in surface water. As can be seen from these studies and the data presented in this paper, *E. coli* resistance to ampicillin is common in the environment. This might be because of leaking wastewater infrastructure. According to a study published in 2022, the number of resistant bacteria found in wastewater was tenfold greater than surface water due to elevated numbers of *E. coli* exposed to high antibiotic concentrations, increasing the chances of resistant genes (Kusi et al., 2022). The leakage of wastewater into surface water increases *E. coli* and antibiotics present in waterways which elevates the number of antibiotic resistant *E. coli*.

The percentage of resistant bacteria found for ciprofloxacin ranged from 0% - 2% (Figure 3b) which is lower than the range of values obtained from Kenyon (2022) and Chen et al. (2017), 22% and 10% respectively. Both studies were conducted in surface water so no comparison could be made with antibiotic resistance in wastewater. However, both studies measured ciprofloxacin concentration in water and its relation to *E. coli* resistance to ciprofloxacin. According to Kenyon (2022), the prevalence of ciprofloxacin resistance in *E. coli* was positively correlated with the increasing concentration of ciprofloxacin in rivers. For instance, the mean ciprofloxacin concentration found in Austria’s rivers was 58.7 ng/L while in China was 329.5 ng/L and *E. coli* resistance to ciprofloxacin was 22% and 56% respectively (Kenyon, 2022). This relationship suggests that the higher the concentration of ciprofloxacin present in waterways, the higher the percentage of *E. coli*’s resistance to ciprofloxacin. Because almost no resistance to ciprofloxacin was found in Waller Creek, this may indicate that there is a low concentration of ciprofloxacin in the creek.

*E. coli*’s resistance to erythromycin in Waller Creek (up to 48%) was greater than the range found by Odonkor & Addo (2018) and Ibekwe et al. (2011) of 23.71% and 21% respectively. Erythromycin is a commonly used antibiotic that is widely utilized to cure skin infections and numerous STDs (MedlinePlus, 2019). One explanation for the higher resistance to erythromycin found in this report may be due to the trend in increasing resistance found over time. *MRSA* resistance to 10 antibiotics was less than 5% in 1999 but increased to between 6% and 18% by 2008, nine years later. Additionally, the usage of these antibiotics to treat *MRSA* increased significantly at the start of the new century, utilizing higher and higher concentrations for better results (Baker et al., 2018). The data from Odonkor & Addo (2018) and Ibekwe et al. (2011) are more than six and twelve years old respectively, thus, resistance might have increased significantly since the previous reports due to continued exposure of *E. coli* in the environment to erythromycin. Alternatively, antibiotic resistance can differ in different locations.

These findings indicate that the concentration of ampicillin and erythromycin are not in a safe range in Waller Creek, increasing the possibility of creating more antibiotic resistant *E. coli* and hence increased health risk to humans. However, without knowing the actual concentration of antibiotics present in Waller Creek (which was beyond the scope of this project), we can only speculate on the low level of resistance to ciprofloxacin compared to ampicillin and erythromycin.

Additionally, it can be noted in Figure 1a that the number of *E. coli* observed October 27, 2023 at Creekside was significantly higher than the number measured during the other collection dates because of the 118.9 mm of precipitation during the four days preceding the collection. Heavy precipitation and discharge increases the number of *E. coli* in urban waters due to stormwater runoff, soil erosion, and untreated sewage directly discharged in urban waterways (Li et al., 2023). The high *E. coli* resistance on October 27, 2023 is correlated with the elevated *E. coli* count measured after heavy precipitation.

Another important result is the high levels of *E. coli* measured on March 12, 2024 at 7^th^ Street compared to the rest of the collection sites collected on the same day (Figure 1a, Figure 2a, and Figure 3a). Rainfall is unlikely to have had a significant impact because it only rained 8 mm four days before. The disproportionate increase of *E coli* collected at 7^th^ Street can be better seen in Figure 2b where the proportion of resistant *E. coli* at 10 μg/mL of erythromycin was 83.97% of the control group with p = 0.4, which indicates no statistical difference. It is likely that the environment around the 7^th^ Street location was a significant factor leading to the high number of antibiotic resistant *E. coli*. Creekside is on the heavily urbanized University of Texas at Austin campus and downstream from St. David’s Hospital, 43^rd^ Street is within a residential area, while 7^th^ Street is in the middle of downtown Austin and has a large homeless population nearby. *E. coli* concentrations near homeless camps can be higher compared to urban waterways that were not near a homeless population (Gerrity et al., 2022). So it may be that the higher levels of *E. coli* and antibiotic resistant *E. coli* are related to the human activities surrounding the 7^th^ Street location.

Among the factors that can influence the levels of antibiotic resistant *E. coli* are location and precipitation. Unsafe antibiotic concentrations and elevated counts of *E. coli* present in urban waterways can increase the transfer of resistance genes between pathogens via horizontal gene transfer (Burmeister, 2015). Since the antibiotic concentrations present in Waller Creek are not known, it is difficult to assess the risk of increasing resistance and how likely it is for horizontal gene transfer to occur.

One limitation of this study was that there was only a single data collection after rainfall, so the results cannot confirm that rainfall increased the quantity of resistant *E. coli* in urban waterways. The same can be said for the increase in *E. coli* seen at 7^th^ Street. Because of budget limitations, collection at different sites could only be done on one day during the collection period, thus there are not replicates of that data to verify if waterways close to homeless camps have a higher percentage of *E. coli* resistant bacteria compared to residential waterways or areas close to hospitals. Further research should be conducted to verify the correlation between precipitation and location with the increase of antibiotic resistant *E. coli*. Despite these limitations, it is clear that Waller Creek has a significant number of antibiotic resistant *E. coli*.

This study found that *E. coli* in Waller Creek have moderate levels of resistance to ampicillin and erythromycin and very little resistance to ciprofloxacin. One explanation is that antibiotics are present in Waller Creek creating a reservoir for antibiotic resistant *E. coli*. Another possibility is that the antibiotic resistant *E. coli* is being introduced from leaking infrastructure or the lack of sanitary resources available to homeless individuals which increases the likelihood of defecation near waterways. For instance, a study conducted in the lower San Diego River Watershed showed that 36% of homeless interviewed practiced outdoor defecation between 4-7 days a week and 6.7% of these individuals stated defecation on low ground near the river (Hinds et al., 2024). Further studies should be conducted in Waller Creek in order to determine the concentration of antibiotics present in water, and its relationship with antibiotic resistant *E. coli*. While there have not been reports of antibiotic resistant infections related to contact with creeks in Austin, TX, the city should monitor the levels of resistance in Waller Creek to determine any changes and take appropriate action.

## Acknowledgments

The authors would also like to express their gratitude to the Green Fund at the University of Texas at Austin for financial support. In addition, they are grateful to IDEXX Laboratories, Inc. for offsetting a portion of the costs of the Colilert. Special thanks to Dr. Ruth Shear for her help with the project and data analysis, and Dr. Timothy Riedel for constructive criticism of the manuscript. Additionally, Emily Payne and Wesley Tran assisted in sample preparation.

